# Exposure to deceased remains of conspecifics extends the lifespan of young and aged *C. elegans* via distinct pathways

**DOI:** 10.64898/2026.02.08.704602

**Authors:** Erik Toraason, Coleen T. Murphy

## Abstract

Signaling factors, both external from an organism’s environment and produced internally by its tissues, regulate the rate of aging. Loss of beneficial signals drives systemic aging, and conversely, restoring these youth-associated signals can rejuvenate an aging individual, as demonstrated by heterochronic parabiosis. Finding factors that promote organismal health and longevity therefore holds great therapeutic promise to slow aging and age-associated disease. Here, we report that exposure to the lysed remains of other worms extends *C. elegans* lifespan. This lifespan extension is not mediated by ascaroside pheromones and is not induced by bacterial cell lysate, suggesting that this effect is not merely produced by nutritional supplementation of cellular contents. We found that a period of discrete exposure at any point across the lifespan is sufficient to induce longevity. However, distinct pathways were activated in young and aged recipients; we found that lysate factors act through insulin/insulin-like growth factor/FOXO signaling (IIS) in young worms, while IIS-independent pathways extend lifespan in older worms. Using fluorescent gene reporter lines, we provide evidence that intestinal IIS is not activated in young worms, suggesting that lysate signals promote longevity via non-intestinal tissues. Our work identifies a novel longevity paradigm in which the remains of deceased *C. elegans* extend the lifespans of living conspecifics through multiple parallel pathways.

## Introduction

Organismal lifespan and aging is regulated by the perception and circulation of factors that promote cellular and systemic integrity. Reintroduction of youth-associated factors can rejuvenate aging organisms, and identifying the specific factors that induce longevity holds therapeutic promise to slow aging and age-associated diseases. Heterochronic parabiosis, the connection of blood circulation between young and old mammals, induces systemic and cognitive rejuvenation in the aged animal^1–6^. Sensory perception of chemicals derived from other organisms can also affect lifespan. In *Drosophila*, aged flies’ lifespans are extended by chemical cues from young flies when they are co-housed^7^. However, the perception of dead conspecifics can inversely shorten lifespan^8,9^. In total, sensory perception and inter-tissue signaling are integrated to inform an organism’s health and lifespan.

The nematode *Caenorhabditis elegans* is a powerful organism with which to dissect mechanisms of aging and rejuvenation^10^. *C. elegans* takes up molecular cues from its environment that directly regulate its physiology and survival; for example, our group discovered that worms interpret small RNAs in pathogenic *Pseudomonas* bacteria to induce gut-to-germline-to-neuron signaling that culminates avoidance of the pathogen^11–16^. This behavior is transgenerationally inherited by the worm’s progeny for four generations^11,15,16^, and worms can further pass this information horizontally via virus-like particles^13^. *C. elegans* also interpret cues secreted from other worms, including dauer pheromone, which affects both developmental decisions and lifespan^25^ ^26^. *C. elegans* sensory perception also impacts its aging; decreasing neuronal activity generally promotes longevity^17^, while neuronal hyperactivation shortens lifespan^18^.

In this study, we test whether *C. elegans* may be used to identify youth-enriched factors that regulate aging by exposing them to the cellular contents of young worms. We hypothesized that this effect could be plausibly mediated by 1) sensory perception of youth or death cues, 2) dietary supplementation of youth-enriched metabolites, or 3) horizontal transfer of pro-longevity signals (‘heterochronic cannibalism’). Using custom microfluidic devices, we indeed observed that exposure to the lysed remains of young *C. elegans* robustly extends recipient worms’ lifespan. This effect is not mediated by known worm pheromones, and bacterial lysate does not induce longevity, suggesting that this intervention is mediated by factors not ubiquitously present in all cell lysates. Strikingly, we find that lysate factors induce longevity via distinct pathways in young and old worms: insulin/insulin-like growth factor signaling (IIS) in young recipient worms, and IIS-independent pathways in aged recipients. In summary, our study defines a novel pro-longevity intervention in *C. elegans*.

## Results

### Exposure to the lysed remains of other worms extends *C. elegans* lifespan

To determine whether exposure to the lysed cellular contents of young *C. elegans* could induce pro-longevity signaling in recipient worms, we used custom microfluidics devices constructed by our group^19^ (Figure 1A) to incubate recipient worms beginning at day 1 of adulthood to the lysed remains of other day 1 adult worms and maintained this intervention through day 14 of the recipients’ adult life (Figure 1B, shaded region is the duration of lysate exposure). We employed these devices to prevent worms from fleeing from the worm lysate on plates, as *C. elegans* can detect and avoid other dead worms^9,20,21^; we later devised a strategy for petri-plate based lysate exposure assays as well (see Methods). Strikingly, exposure to day 1 adult wild-type lysate increased *C. elegans* recipients’ mean lifespan by 21% (p<0.001; Figure 1B).

**Figure 1.**
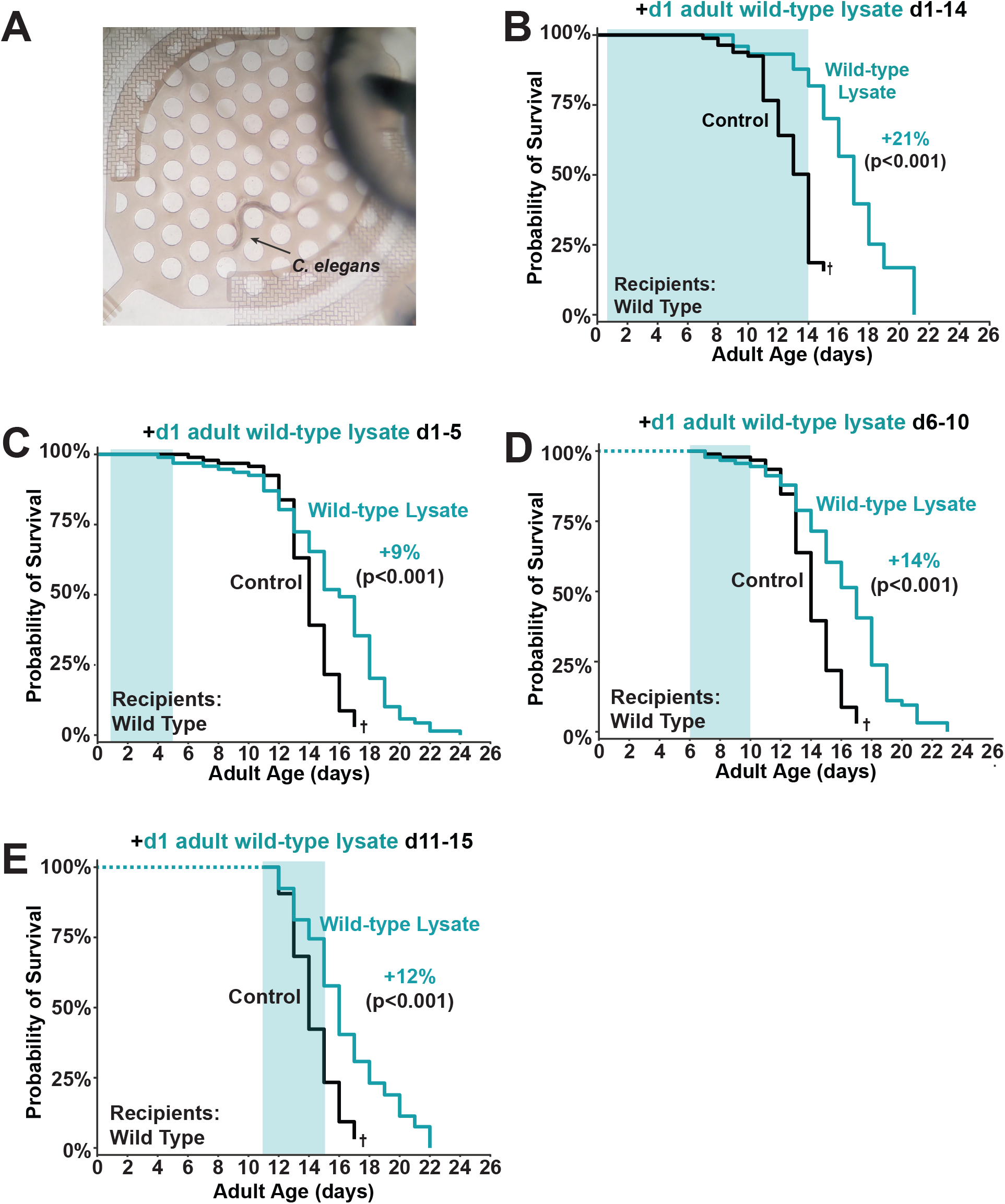
Exposure to *C. elegans* lysate extends recipient worms’ lifespan. A) Image of a single well of a CeLab chip^19^ containing an N2 wild-type *C. elegans* worm. All experiments in this figure were performed using the CeLab chip. B) Exposure to 5μm filtered wild-type *C. elegans* lysate for the first 14 days of adult life extends lifespan (1 representative replicate of 2). C-E) Exposure to 5μm filtered wild-type *C. elegans* lysate in early life (C), mid life (D), or late life (E) extends lifespan. P values in B-E were calculated with the Log-Rank test. C-E were analyzed using the same control population.

Multiple interventions that extend *C. elegans* lifespan also slow reproductive aging^19,22–24^. To test whether lysate exposure delayed oocyte aging, we examined the reproductive span of hermaphrodites exposed to *C. elegans* lysate. We found that mated *C. elegans*, for which sperm is not limiting and oocyte decline drives reproductive cessation, did not exhibit an altered reproductive span (Supplemental Figure 1A), and self-mated *C. elegans* hermaphrodites exhibited a slight (5.5%; p=0.02) increase in mean reproductive span (Supplemental Figure 1B). As self-mated *C. elegans* exhaust their self sperm supply before their oocytes exhibit meaningful decline, this phenotype suggests that exposure to *C. elegans* lysate may very slightly slow ovulation in unmated worms. Thus, while lysate exposure significantly increases somatic lifespan, it does not proportionally slow *C. elegans*’ reproductive aging.

Transfer of biomolecules from young to aged individuals in heterochronic parabiosis paradigms rejuvenates aging organisms in part by increasing the abundance of youth-enriched factors in aged individuals^1–4^. To test whether exposure to *C. elegans* lysate extended lifespan by inducing lifestage-specific signals, we incubated *C. elegans* with lysate from young (day 1 adult) wild-type worms in discrete windows in the recipients’ early life (adult day 1-5, Figure 1C), mid life (adult day 6-10, Figure 1D), or late life (adult day 11-15, Figure 1E). We found that lysate exposure at any of these timepoints was sufficient to extend lifespan (Figure 1C-E). Thus, our data demonstrate that *C. elegans* lysate factors can induce longevity across a recipient’s lifespan.

### Lysate factor-mediated longevity is not mediated by ascaroside pheromones

*C. elegans* synthesizes and secretes ascaroside pheromones, which serve as signals for mating attraction and as proxies of population density and thus potential food scarcity^25^. Exposure to high concentrations of ascarosides also extends *C. elegans*’ lifespan^26^. As worm lysates contain ascarosides, we reasoned that the lifespan extension that we observed may be mediated by pheromones. However, we observed that lysate from *daf-22* mutants^27^, which are defective in ascaroside biosynthesis, robustly extended lifespan when applied to young or to aged recipient worms (Figure 2A-B). Thus, the primary lifespan-extending factor(s) in worm lysate is not a canonical ascaroside.

**Figure 2.**
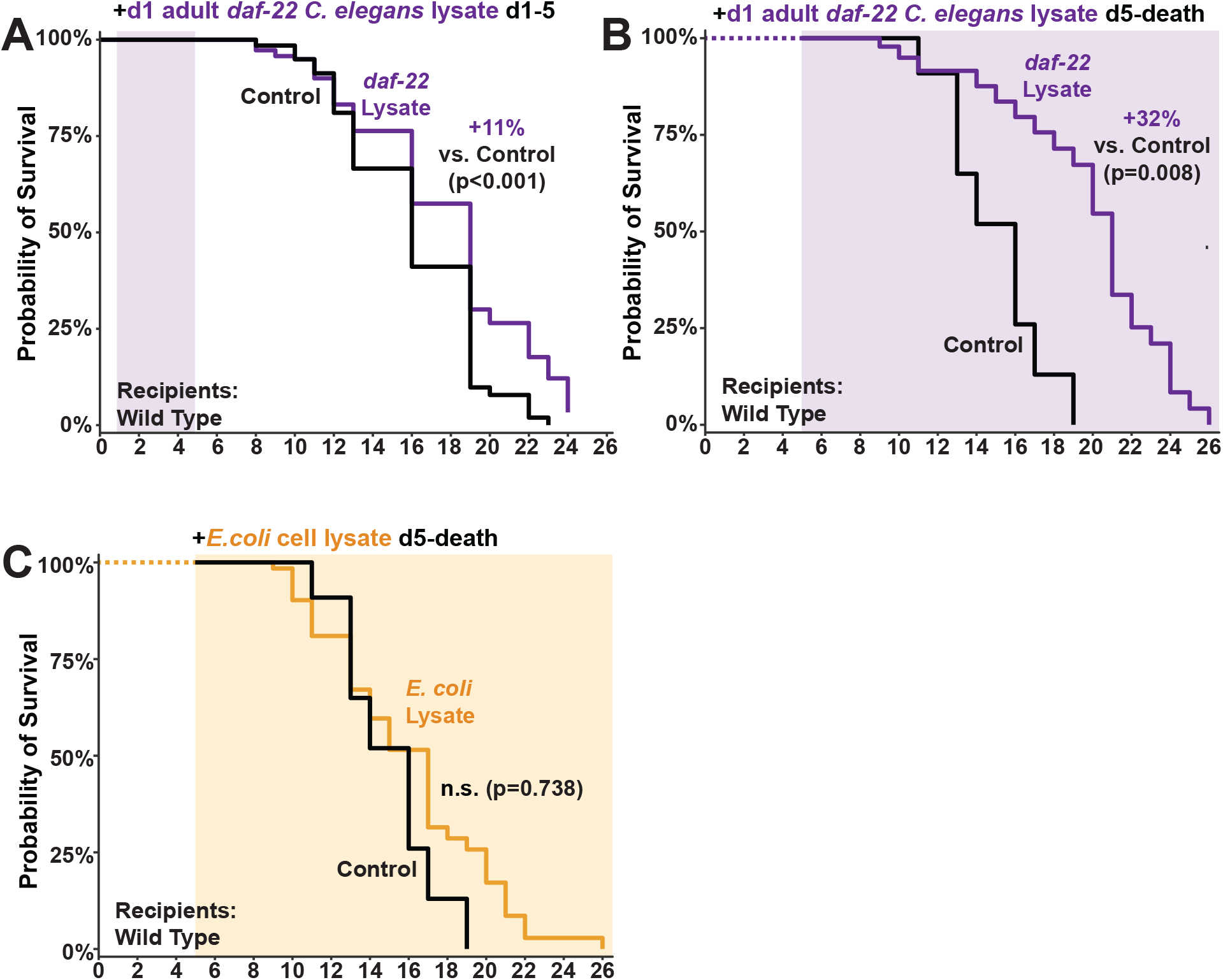
Lifespan-extending factors in *C. elegans* lysate are not ascarosides and are not ubiquitously present in all cell lysates. A-B) Exposure to 0.2μm filtered lysate from day 1 adult *daf-22 C. elegans* ascaroside biosynthesis mutants extends lifespan both in early (A) and late (B) life. C) Lysate from OP50 *E. coli* does not extend lifespan. P values in all panels were calculated with the Log-Rank test.

### Bacterial cell lysate does not extend *C. elegans* lifespan

We next asked whether the components of *C. elegans* lysate that extend lifespan were ubiquitously present in all cell lysates by exposing worms to lysate from *Escherichia coli* OP50 bacteria (Figure 2C); however, *E. coli* lysate did not extend *C. elegans’* lifespan (Figure 2C). Thus, we concluded that the longevity-promoting factors were not ubiquitously present in all cell lysates. Importantly, this result also suggests that lifespan extension by worm lysate exposure was not merely a product of supplying “pre-digested” nutrients to worms and was instead a product of a more specific signaling or dietary mechanism.

### *C. elegans* lysate exposure in early life extends lifespan via insulin/insulin-like growth factor signaling

Decreased insulin/insulin-like growth factor signaling (IIS) extends lifespan across phyla in worms^10,28^, flies^29,30^, and mice^31^. In *C. elegans*, reduced signaling from the DAF-2 insulin receptor induces translocation of the DAF-16/FOXO transcription factor into the nucleus, where it induces the transcription of genes that increase cellular stress resilience and longevity^32^. To determine if lysate exposure extends lifespan via IIS/FOXO, we exposed *daf-16* null mutants to wild-type *C. elegans* lysate in early life (adult day 1-5, Figure 3A) or mid-late life (adult day 6-10, Figure 3B). Strikingly, young *daf-16* mutants did not exhibit lifespan extension (Figure 3A), but aged *daf-16* mutant mean lifespan was increased by >20% by this intervention (Figure 3B).

**Figure 3.**
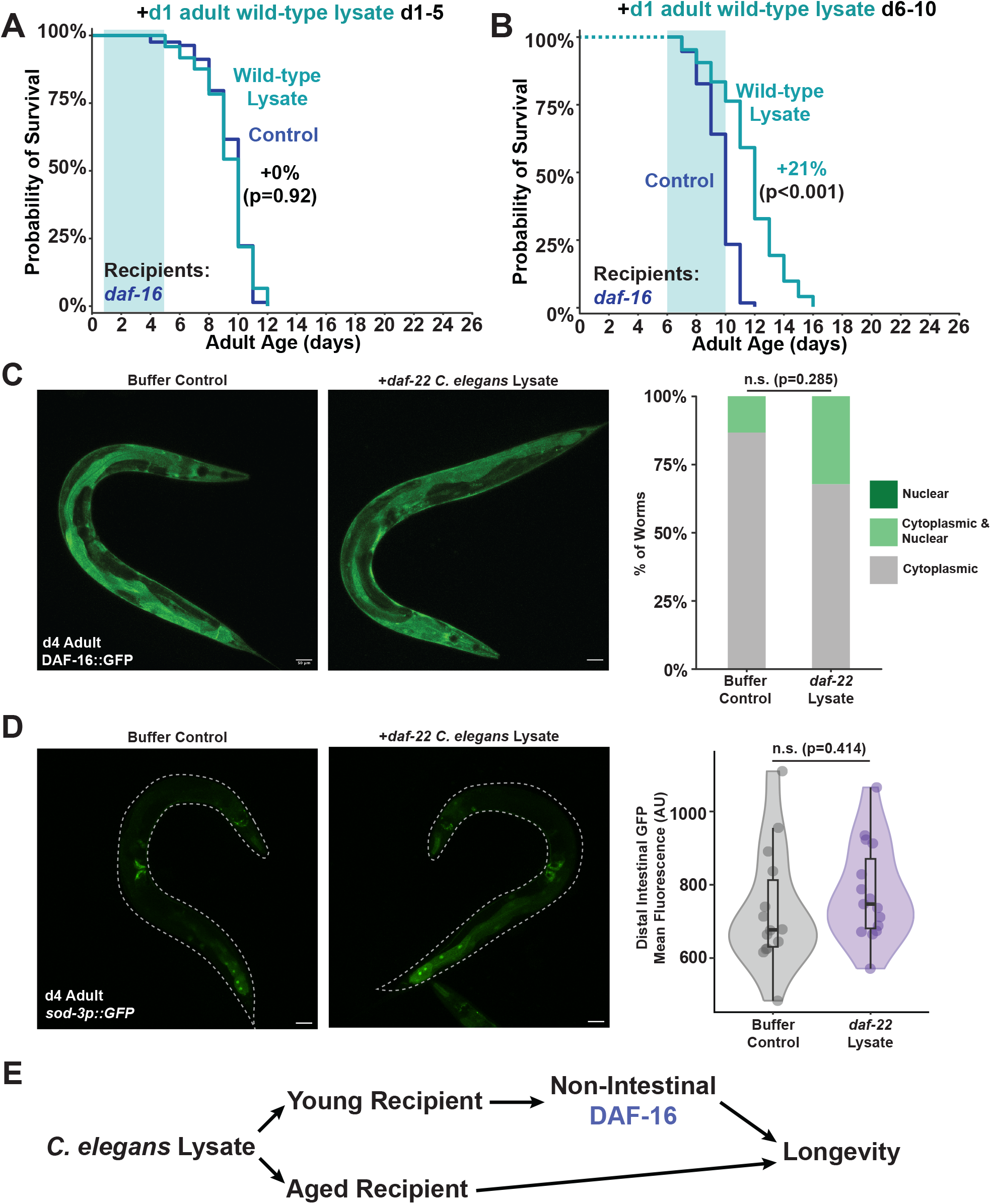
Lysate exposure in early life, but not late life, extends longevity via insulin/insulin-like growth factor signaling. A-B) Exposure of *daf-16* recipient worms to 5μm filtered wild-type *C. elegans* lysate extends lifespan in early life (A) but not in mid-late life (B). P values in A-B were calculated with the Log-Rank test. C) Images and quantification of DAF-16::GFP localization in the intestine of day 4 adult worms exposed to 0.2μm filtered lysate from day 1 adult *daf-22 C. elegans* ascaroside biosynthesis mutants (exposed from adult day 1-4). P value was calculated by Chi Square test. D) Images and quantification of *sod-3p::GFP* fluorescence in the distal intestine (see Methods). In microscopy images, worms are outlined with dashed lines. P value was calculated by Student”s t-test. In C and D, scale bars represent 50μm. E) Proposed pathway by which lysate factors extend recipient worms” lifespan. In panels with box plots, the upper and lower hinges indicate the first and third quartiles, while the whiskers indicate the minima and maxima (interquartile range ±1.5*IQR), and horizontal lines indicate the median. A-B were analyzed using the same control population.

The majority of IIS contributions to lifespan are mediated by DAF-16 activity in the intestine^33,34^. We therefore hypothesized that lysate exposure-induced longevity in young *C. elegans* was mediated by intestinal IIS. When DAF-16 is activated to induce gene transcription, it translocates from the cytoplasm to the nucleus. However, we did not observe significantly increased nuclear translocation of DAF-16 in young worms exposed to *C. elegans* lysate (Figure 3C). We similarly did not observe significantly increased expression of the intestinal IIS reporter *sod-3* Figure 4D). Together, these reporter line results suggest that lysate factors activate IIS/FOXO in a non-intestinal tissue to promote longevity.

In total, our data support a model in which factors in *C. elegans* lysate act on distinct pathways in young and aged recipient worms to regulate aging (Figure 3E). We propose that in young worms, lysate factors activate non-intestinal IIS to extend lifespan, while in aged worms lysate factors induce IIS-independent mechanisms of longevity.

## Discussion

Our study addresses a fundamental question: how does encountering and/or consuming the cellular components of a healthy cell impact another organism’s aging? Using *C. elegans*, we demonstrated that exposure to the lysed remains of young adult worms extends recipients’ lifespan. This effect is not mediated through known ascaroside pathways, and also is not induced by bacterial cell lysate. Although *C. elegans* recipients of any age can benefit from this intervention, we found that lysate factors act on distinct pathways depending on the recipients’ age: IIS-dependent pathways in young animals (but not intestinal IIS) and IIS-independent pathways in aged worms.

While our work shows that exposure to *C. elegans* lysate can extend lifespan, Hernandez-Lima et al. (2025)^9^ observed that incubating *C. elegans* with the corpses or lysed remains of sodium-azide-killed worms reduces lifespan^9^. One notable difference between our approaches is that we lysed worms while they were still living (see Methods); it is possible that killing worms in 1M sodium azide (as performed by Hernandez-Lima et al.) depletes pro-longevity molecules or induces the production of lifespan-shortening compounds that overrides them. Thus, the mode of lysate preparation is likely important to determine whether lysate exposure extends or decreases *C. elegans* lifespan.

In a parallel study (Toraason et al. *bioRxiv* 2026)^21^ we mapped the neuronal mechanisms of *C. elegans* necrotaxis (death avoidance) to a novel small-molecule cue that we named ‘Todstoff’. That *C. elegans* flees other dead worms when factors in dead worms extend their lifespan raises the possibility that the lifespan-extending pathways that are induced by exposure to lysate are a byproduct of stress signaling or hormesis. Under this model, worms may enact hierarchical responses when they encounter the remains of another worm: their first strategy is to necrotax away to avoid danger; however, if they cannot escape, then they may upregulate stress response pathways to protect themselves from whatever threats killed their kin.

Aside from encountering an injured or killed adult worm, the other context in which *C. elegans* might interact with the cellular contents of another worm is if they hatch inside their mother. A subpopulation of *C. elegans* hermaphrodites undergo matricidal hatching (‘bag-of-worms’ phenotype)^35^, when they fail to lay eggs and their offspring hatch inside of them and consume them as food. Matricidal hatching increases during starvation^36^ and as worm age^35^. While our study focused on the responses of adult worms to cell lysates, it is possible that some of these longevity pathways (particularly the IIS-dependent pathway in young adult recipients) are shared with larval animals. Thus, some longevity-promoting lysate cues may in fact be altruistic signals that are interpreted by the hatched progeny of dying mothers to instill resilience in the offspring. In total, our work uncovers novel modalities of lifespan extension in *C. elegans*.

## Methods

### *C. elegans* culture and maintenance

*C. elegans* strains were maintained at 20°C under standard conditions on high growth medium (HG) agar plates [3g/L NaCl, 20g/L Bacto-peptone, 30g/L Bacto-agar, 4mL/L cholesterol (5 mg/mL in ethanol), 1 mL/L 1M CaCl_2_, 1 mL/L MgSO_4_, and 25mL/L 1M KPO_4_ pH6.0] seeded with OP50 *Escherichia coli* bacteria. Worm populations were synchronized by bleaching in alkaline-bleach solution [250mM KOH, 12% hypochlorite bleach] followed by two washes with M9 buffer [6 g/L Na_2_HPO_4_, 3g/L KH_2_PO_4_, 5 g/L NaCl, 1 mL/L 1M MgSO_4_ in ddH2O].

Strains used in this study include:

N2 (wild type)

CF1038 *daf-16(mu86)* I

CF1553 muIs84 [(pAD76) *sod-3p::GFP* + *rol-6(su1006)*]

DR476 *daf-22(m130)* II

TJ356 zIs356 [*daf-16p::daf-16a/b::GFP* + *rol-6(su1006)*]

### Preparation of lysates

Bleach synchronized worms were raised on HG plates seeded with OP50. Day 1 adults were collected by washing with PBS buffer pH 7.4 [0.137M NaCl, 2.7mM KCl, 8mM Na2HPO4, 2mM KH2PO4] (Invitrogen) if lysates were to be used for plate assays or S Medium (for 1L: 5.85 g NaCl, 1 g K2 HPO4, 6 g KH2PO4, 1 ml cholesterol (5 mg/ml in ethanol), 10 ml 1 M potassium citrate pH 6, 10 ml trace metals solution, 3 ml 1 M CaCl2, 3 ml 1 M MgSO4) if they were to be used in CeLab chips. Trace metals solution contained (in 1L) 1.86 g disodium EDTA, 0.69 g FeSO4…7 H2O, 0.2 g MnCl2…4 H2O, 0.29 g ZnSO4…7 H2O, 0.025 g CuSO4…5 H2O. Worms were washed 3x with PBS or S Medium to remove residual bacteria. Worms were then pelleted by gravity on ice and excess buffer was removed. Worms were then lysed on ice by Dounce homogenization or gentle sonication using a needle sonicator. The lysate was then centrifuged at 500xg for 10 minutes in a 4°C refrigerated benchtop centrifuge to separate carcasses from lysate, after which the supernatant was collected and filtered with a 5μm or 0.2μm syringe filter (see figure legends for specific experiments). Protein content of the lysate was used as a proxy for lysate concentration across trials, as determined using a Qubit Protein Assay kit (ThermoFisher). Lysate was either immediately used in experiments or was flash frozen in liquid nitrogen and stored at-80°C for later use.

### Lifespan assays in CeLab microfluidic chips

Worms to be used in lifespan assays were bleached to NGM plates (3 g NaCl, 17 g agar, and 2.5 g peptone in 1L H2O) seeded with OP50 *E. coli* bacteria before loading into CeLab chips. CeLab chips were used as described in Sohrabi et al. 2023^19^. Worms in CeLab chips were fed heat-inactivated OP50 as described in Sohrabi et al. 2023. Worms were scored as dead when they did not respond to gentle stimulation from the pins in the microfluidic chip wells. Worms that bagged or exploded in the were censored. Control chips were loaded with additiona S medium (negative buffer control) while lysate was loaded at a final 500μg/μL normalized protein concentration. For lifespans in which worms were exposed to lysate later in life (d5+), worms that died before the exposure began were censored.

### Reproductive span assays in CeLab microfluidic chips

Reproductive spans of mated and unmated worms were performed as described in Sohrabi et al. 2023^19^.

### Lifespan assays on plates

Plate lifespan assays were performed using 35mm petri plates containing NGM agar supplemented with 300μg/mL streptomycin seeded with OP50-1 (streptomycin resistant OP50). On the day before worms were added, plates were administered with 100μL of PBS (negative buffer control) or worm lysate in PBS (5mg/mL protein content).

### Microscopy and image quantification

Worms were maintained as per lifespan assays on plates (see above) and were exposed to *daf-22 C. elegans* lysate on adult days 1-4. On the day of imaging, day 4 adult worms were picked onto an agar pad with a 10μL drop of M9 with levamisole to paralyze them and were imaged within 30 minutes. Images were acquired using a Nikon Eclipse Ti microscope at 20x magnification. Images were analyzed using Fiji. For quantification of DAF-16::GFP, images were blinded before scoring. *sod-3p::GFP* was quantified from maximum intensity projections (FIJI) from an ROI tracing the posterior intestine (from vulva to tail) to find the mean fluorescence intensity.

### Statistics

All statistics were performed using R (v4.4.2). Data wrangling was performed using the reshape2 (v1.4.4), DescTools (v0.99.53), and tidyverse (v2.0.0) packages. Lifespan analysis was performed using the ggsurvfit (v1.1.0), survival (v3.6-4), survminer (v0.4.9), and gtsummary (v2.0.3) packages. Specific statistical tests used are denoted in figure legends.

## Acknowledgements

Funding for this work was provided by the Simons Foundation Collaboration on Plasticity in the Aging Brain (SCPAB) to CTM, an NIH Director’s Pioneer Award to CTM (DP1AG077430), and the Glenn Foundation for Medical Research. ET is a Simons SFARI Awardee of the Life Sciences Research Foundation postdoctoral fellowship. Some strains were provided by the *Caenorhabditis* Genetics Center (CGC), which is funded by NIH Office of Research Infrastructure Programs (P40 OD010440).

---

**Supplementary Figure 1.**
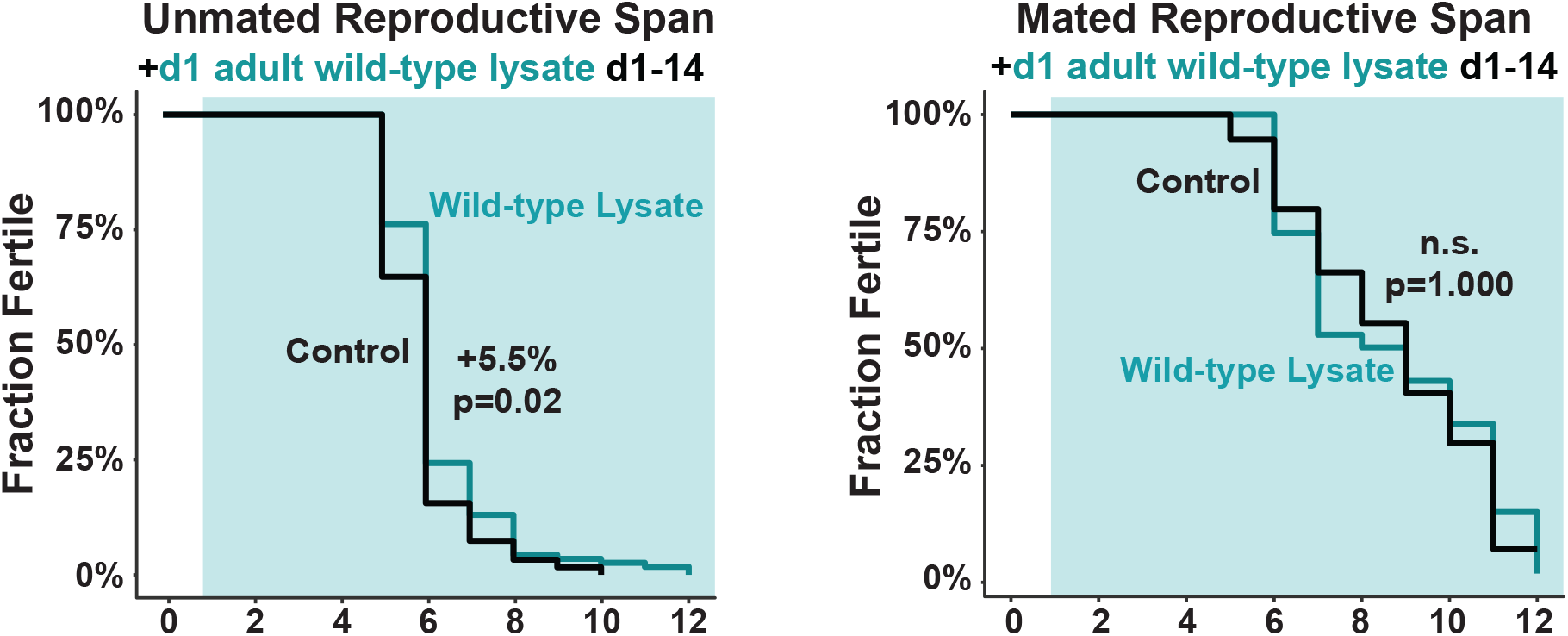
Reproductive span assays performed in CeLab chips using mated (above) and unmated (below) N2 wild-type worms. P values were calculated by Log-Rank test.

